# Virtual reality headset geometry constrains dorsolateral prefrontal cortex targeting with transcranial magnetic stimulation

**DOI:** 10.64898/2026.08.11.744141

**Authors:** Franka Arden, Phil Henneken, Zsolt Turi, Andreas Vlachos

## Abstract

**Background:** The integration of virtual reality (VR) and non-invasive brain stimulation (NIBS), particularly transcranial magnetic stimulation (TMS), represents a promising approach for closed-loop neuromodulation. Yet the concurrent application remains limited, partly due to insufficient characterization of hardware compatibility of head-mounted displays with standard TMS coil placement protocols.

**Objective:** To systematically quantify the coil-to-scalp distance constraints imposed by VR headsets across cortical targets and coil orientations and to determine feasible intensity compensation ranges based on stimulator output parameters.

**Methods:** Neuronavigated coil positioning was performed on five anatomically realistic 3D-printed head models across 26 scalp positions in eight coil orientations based on the 10-10 EEG system and dorsolateral prefrontal cortex (DLPFC) using two VR headsets of notably different form factors (Meta Quest 2 and Bigscreen Beyond). The deviations of coil positions from intended targets were registered and quantified as coil-to-scalp distance displacement. Individual electric field (E-field) simulations were conducted in SimNIBS at the F3 position across 4-40 mm coil-to-scalp distance to characterize field decay and assess the limits of intensity compensation.

**Results:** Both in the directed DLPFC targeting and in systematic scalp positions evaluation, the Meta Quest 2 headset substantially increased coil-to-scalp distance over prefrontal regions, exceeding the compensable range across all metrics. The Bigscreen Beyond headset produced significantly smaller coil-to-scalp distance displacement in prefrontal regions, remaining within feasible E-field intensity compensation limits. Single-pulse and iTBS protocols did not induce functional interference with the hardware under realistic targeting conditions.

**Conclusion:** VR headset geometry is the primary determinant of concurrent VR-TMS feasibility. The findings define practical quantitative hardware design requirements and boundaries for future integrated VR-TMS systems and provide a practical framework for optimizing existing VR-TMS protocols.

## INTRODUCTION

Fully immersive virtual reality (VR) and transcranial magnetic stimulation (TMS) have each independently advanced the fields of neuroscience and neuromodulation, yet their physical integration remains an engineering challenge. Head-mounted VR displays occupy the cranial region critical for TMS coil placement, potentially introducing spatial interference of the devices and coil-to-scalp distance constraints, which could compromise stimulation efficacy. Despite growing interest, this hardware compatibility problem has not been characterized. The combined use of these technologies has demonstrated value in basic research on the neural basis of cognition and behavior, as well as clinical applications (Buch et al., 2011; Sofroniew et al., 2014; Connelly et al., 2024; Gainsford et al., 2020; Li & Zanto, 2024; Philip et al., 2024; van’t Wout-Frank et al., 2024; Lefaucheur et al., 2017), but despite this increase in their reported use, fully immersive closed-loop VR-TMS paradigms have not been systematically pursued, neither technically nor clinically.

Emerging evidence suggests that combining TMS with both semi-immersive and fully immersive VR can enhance neurophysiological and behavioral effects, underscoring the clinical potential of this approach (Mishra et al., 2021; Chauhan et al., 2024; Afonso et al., 2024; Teo et al., 2016). However, most existing studies have employed asynchronous designs or non-immersive setups, limiting the temporal alignment between stimulation and behavioral states. Several methodological approaches have been proposed to address timing and synchronization issues (Elze, 2010; Garaizar et al., 2014; Wiesing et al., 2020; Bridges et al., 2020), yet, fully integrated, concurrent VR-TMS applications remain rare. When concurrent approaches have been implemented, they have primarily targeted the motor cortex (Bagce et al., 2012; Shin et al., 2024) or used alternative stimulation modalities such as transcranial direct current stimulation (tDCS) (Kim Y. J. et al., 2014; Massetti et al., 2017). Consequently, critical questions remain regarding the feasibility and constraints of targeting non-motor regions, particularly in prefrontal areas, using VR headsets.

TMS is an established method for the diagnosis and treatment of neurological and psychiatric disorders (Lefaucheur et al., 2020), with approval from the United States Food and Drug Administration (FDA) for treating major depressive disorder (Janicak et al., 2008; O’Reardon et al., 2007; Carmi et al., 2019), obsessive-compulsive disorder (Carmi et al., 2019), migraine-associated pain (Lipton & Pearlman, 2010), and smoking cessation (Zangen et al., 2021; Harmelech et al., 2023). It is widely used in clinical neurophysiology and cognitive neuroscience and has increasingly been integrated into multimodal and closed-loop paradigms. These include combinations with electroencephalography (EEG; Hernandez-Pavon et al., 2023), magnetic resonance imaging (MRI; Bergmann et al., 2021), near-infrared spectroscopy (NIRS; Curtin et al., 2019), and virtual reality (VR; Banduni et al., 2023) to probe mechanisms of plasticity, connectivity, and higher-order cognitive functions such as learning and memory (Diester et al., 2024). Collectively, these approaches highlight the capacity of TMS to modulate neural circuits and support targeted, mechanistically informed interventions.

In parallel, VR has gained considerable interest in clinical contexts, with applications spanning stroke and brain injury rehabilitation, phobias, post-traumatic stress disorder, cerebral palsy, chronic pain, anxiety and depressive disorders, addiction and neurodegenerative conditions (Chen et al., 2022; Spreij et al., 2014; Rothbaum et al., 1995; Shiban et al., 2017; Kothgassner et al., 2019; Liu et al., 2022; Matamala-Gomez et al., 2019; Zeng et al., 2018; Bouchard et al., 2017; Mazza et al., 2021; Ferrer-García et al., 2017; Cheng et al., 2022; Doniger et al., 2018; Ford et al., 2023). The integration of immersive VR with NIBS is particularly promising, as it may enhance experimental control and enable temporally precise, task-dependent neuromodulation, including in closed-loop settings (Cassani et al., 2020; Drigas & Sideraki, 2024; Stramba-Badiale et al., 2020; Weber et al., 2021). However, most studies to date have relied on semi-immersive, screen-based VR paradigms rather than fully immersive systems. With rapid advances in VR technology, the level of immersion has emerged as an important experimental and clinical variable. Fully immersive approaches, such as head-mounted displays or Cave Automatic Virtual Environments, can reduce external sensory input and improve control over behavioral context, thereby enabling more precise engagement of defined neural networks (Geraets et al., 2021; Freeman et al., 2017; Cybinski et al., 2024; Kim K. A. & Ahn, 2024).

In this study we provide the first systematic characterization of the VR-TMS compatibility constraints. Using neuronavigated coil positioning on anatomically realistic 3D-printed head models, we quantified coil-to-scalp distance displacements in dorsolateral prefrontal cortex (DLPFC) defined by MNI coordinates and the 10-10 EEG system, followed by a systematic assessment of 26 10-10 EEG system scalp positions in eight coil orientations. The extent of E-field decay and its compensation through stimulation intensity scaling indicated that E-field reductions can be compensated within defined limits. Our findings show that the headset form factor, rather than stimulation adjustment, determines the feasibility of concurrent prefrontal applications, and define compensation thresholds that can help navigate the development of hardware-compatible immersive VR-TMS systems.

## MATERIALS AND METHODS

### Ethics

MRI data were obtained from original studies, ethics statement was reviewed and approved by the Ethics Committee of the University Medical Center Göttingen (Application number: 35/7/17; Zmeykina et al., 2020) and University of Freiburg (Application number: 23-1114-S1). Written informed consent was obtained from all participants. One dataset was acquired from a public resource (Lynch et al., 2023). No MRI data has been collected for the present study, all datasets were obtained from previous studies or public resources.

### Experimental Design

Five anatomically realistic 3D-printed head models were generated from MRI data using the open-access software package for the Simulation of Non-invasive Brain Stimulation (v. 3.2.6; v. 4.0.0; v. 4.5) (SimNIBS; Thielscher et al., 2015). The accessibility of the left DLPFC was assessed using the 5 cm rule (George et al., 1995), targeting a predefined left M1 position in MNI/RAS space (approximately −50.6; −17.0; 59.0) and M1 via anatomical landmarks (DaSilva et al., 2011). Anatomical landmark for left M1 was defined as 20% of the distance from vertex to the left pre-auricular point, while the vertex was calculated by identifying the cross-point of the halfway between inion and nasion and the distance between pre-auricular points. Additionally, we used the F3 position of the 10-10 EEG system, corresponding to DLPFC (Herwig et al., 2003). Coil positions were registered with a neuronavigation system (Localite TMS navigator, Localite GmbH, Bonn, Germany), each measurement was repeated five times with and without the VR headset. To systematically assess brain region accessibility, 26 positions from the 10-10 EEG system were evaluated on the left hemisphere. The 10-10 EEG coordinates were obtained via SimNIBS. Within each position, 8 coil orientations were evaluated, by rotating the coil 45 ° around the Z-axis (Localite). While anatomical variability may lead to slight differences in spatial direction between positions, orientations are comparable across subjects. Coil positioning accuracy was ensured using neuronavigation and any interference from the headset was managed by adjusting the coil-to-scalp distance (X-axis in Localite system, corresponding to the Z-axis in SimNIBS). E-field simulations were performed using SimNIBS and safety evaluations were conducted with the coil positioned directly touching and at 3.5 cm above the VR headset.

### Head Model Generation and 3D printing

Head models were generated from T_1_-weighted and T_1_/T_2_ weighted MRI images using SimNIBS. Either ‘charm’ (SimNIBS 4.0.0, n=2) or ‘headreco’ (SimNIBS 3.2.6, n=3) functions were employed, reflecting the software version availability during the study timeline. All head models, the TMS coil (Cool-B65, MagVenture, Denmark), and the Bigscreen Beyond VR headset (standard factory dimensions) were 3D-printed using a Raise3D Pro3 printer with Hyper Speed Kit (PLA 1.75 mm; 0.4 mm nozzle; 0.2 mm layer height; 2.0 shells; 5% infill density; touch platform only support).

### VR Headsets

A Meta Quest 2 VR headset (Meta Platforms, Inc., USA) was used in this study. To evaluate the impact of a smaller VR headset (Table 1), a 3D-printed model of the Bigscreen Beyond VR headset (Bigscreen Inc., USA) was tested, as the headset was not yet commercially available at the start of the project (available at https://sketchfab.com/ (Official 3D Printable Model)). To validate the geometric accuracy of the printed model, additional control measurements were performed using the actual Bigscreen Beyond headset in one representative subject across seven prefrontal positions. No substantial deviations were observed.

**Table 1:** Dimensions of popular VR headsets.

| Headset | Length (mm) | Width (mm) | Height (mm) |
| --- | --- | --- | --- |
| Apple Vision Pro | 248 | 89 | 159 |
| Bigscreen Beyond (without strap) | 52 | 143 | 24 |
| Bigscreen Beyond 2 (without strap) | 52 | 143 | 24 |
| HP Reverb G2 | 186 | 75 | 84 |
| HTC Vive (without strap) | 190 | 117 | 140 |
| Meta Quest 2 (without strap) | 191.5 | 102 | 142.5 |
| Meta Quest 3 (without strap) | 184 | 98 | 160 |
| Meta Quest 3s (without strap) | 184 | 98 | 160 |
| Pico 4 (with strap adjustments) | 255-310 | 163 | 80 |
| Sony PS VR2 | 278 | 212 | 158 |
| Valve Index (with strap adjustments) | 290-310 | 260 | 110 |

### Electric Field Simulations and Analysis

E-field simulations were conducted in SimNIBS. Stimulation intensity was set at a coil-current rate of 1.49×10^6 A/s, equivalent to 1% maximum stimulator output (MSO; MagPro X100 stimulator, Cool-B65 coil, MagVenture, Denmark). A linear relationship between MSO and the electric field was assumed. The reference simulations were performed at the F3 position with a coil-to-scalp distance of 4 mm. A single standardized anterior-posterior coil orientation was selected to isolate distance effect from orientation-dependent variability. For clinical interpretation, we used published values indicating that 100% of the resting motor threshold (RMT) for the MagVenture Cool-B65 coil corresponds to approximately 51.7% MSO. In therapeutic practice, intensities usually range between 80 and 120% RMT, corresponding to 41-62% MSO. Thus, 62% MSO was taken as the upper end of routinely applied stimulation, and the theoretical ceiling of 100% MSO defined the maximum feasible scaling window.

The region of interest (ROI) was defined once from the 4 mm baseline simulation as a 10 mm diameter sphere constrained to gray matter, centered at the cortical voxel exhibiting peak field intensity in the 4 mm baseline simulation. This ROI mask was then applied consistently across all subsequent simulations to enable direct comparison of field parameters across distances. Within the ROI, we extracted field characteristics including maximum, mean, median, and the 98th percentile (P98). Focality was assessed as a metric of GM volume with an electric field magnitude exceeding 75% or 50% of the 99.9th percentile peak value. To evaluate the effect of increased coil-to-scalp distance on field characteristics and stimulation feasibility, simulations were performed at 1 mm increments from 4 to 40 mm, with all other parameters held constant. The scaling factor required to restore the reference field parameters was calculated for each metric as the ratio of the reference value to the value obtained at each distance.

### Technical Compatibility

The evaluation of technical compatibility involved positioning the coil either directly on top of the VR headset or 3.5 cm above it during stimulation, as well as mounting the headset on a head model with the coil targeting the F3 location. To assess potential interference between the stimulation and the image and sound quality of the VR headset, all experiments were conducted while playing a high-quality video with dynamic sound and movement (https://www.youtube.com/watch?v=ftp_gXL0NlU). Video and audio output from the VR headset were recorded using a Sony Alpha 6700 4K digital camera (Sony, Japan) during stimulation.

### Data analysis, statistics, and digital illustrations

Data analysis and visualization were performed in Python (v. 3.11.3; Python Software Foundation) using the NumPy, pandas, NiBabel, SciPy and Matplotlib libraries. Differences in coil-to-scalp distance displacement between headsets at each coil orientation were assessed using paired samples t-tests. Normality of difference distributions was confirmed using the Shapiro-Wilk test (p > 0.05 for all comparisons). Data organization was performed in Microsoft Excel (Microsoft Corp., USA). Figures were prepared using Matplotlib, BioRender (BioRender, USA), Inkscape (Inkscape Project), and Microsoft PowerPoint (Microsoft Corp., USA). Code architecture, troubleshooting and error handling were supported in part by AI-assisted programming tools (Claude, Anthropic). All scripts were reviewed, tested and validated by the authors and are available upon reasonable request.

## RESULTS

### Frontal cortex targeting constraints with a VR headset

We assessed frontal cortex targeting using five individual head models (Figure 1A). For both primary motor cortex (M1) and DLPFC targets, the 3D-printed coil was positioned tangentially to the head surface, with the handle oriented approximately 45° to the sagittal plane.

**Figure 1:**
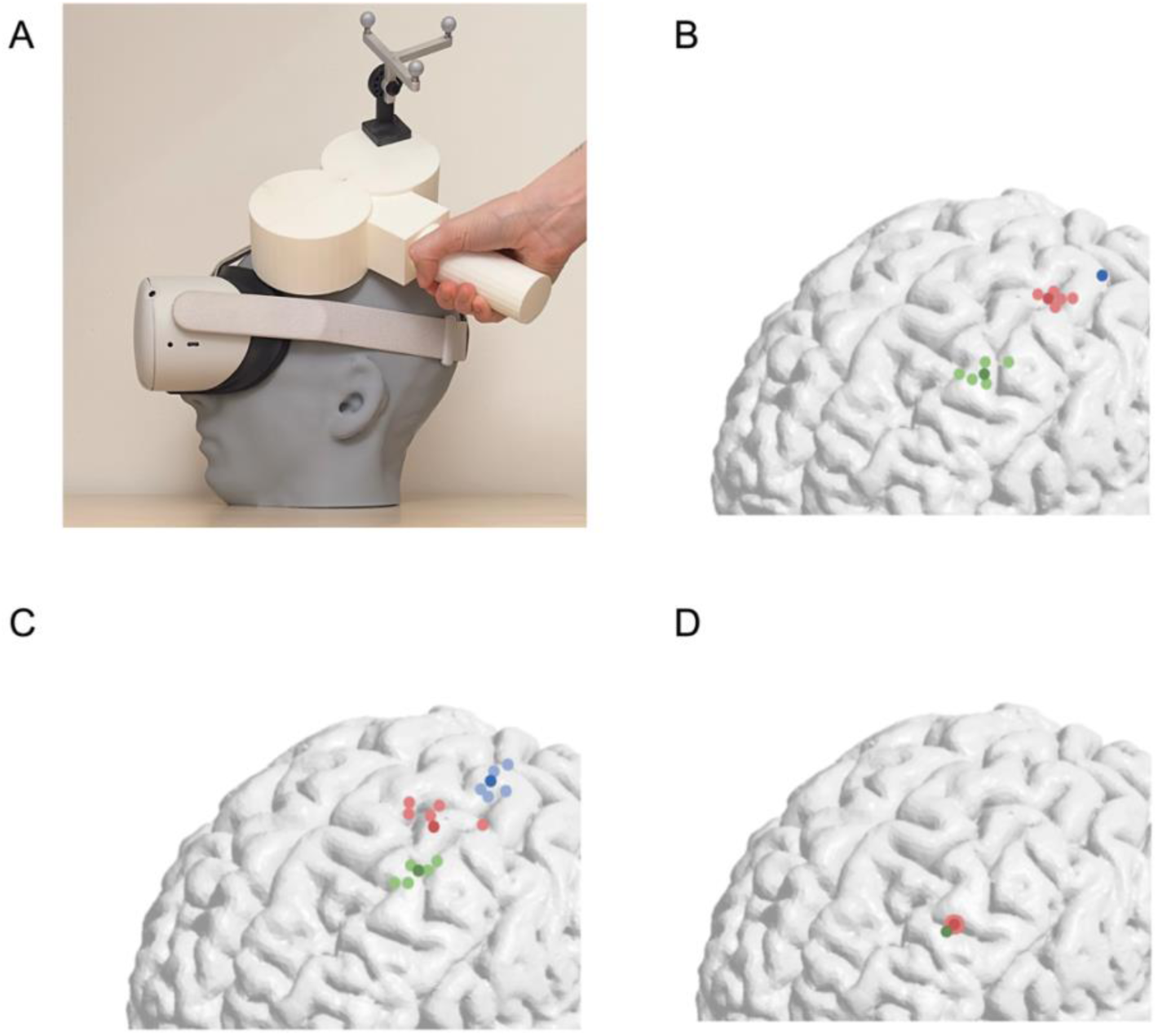
Targeting the primary motor cortex (M1) and dorsolateral prefrontal cortex (DLPFC). **(A)** Experimental setup, illustrating the 3D-printed head model, figure-of-eight coil and Meta Quest 2 VR-goggles. One of five head models is shown; comparable results were obtained across all models (Figure 2). **(B)** Coil center positions based on M1 MNI coordinates and the 5-cm method for targeting DLPFC. M1 (blue), DLPFC without VR headset (light green; mean in dark green), and DLPFC with VR headset (light pink; mean in dark pink). **(C)** Coil center positions determined using anatomical landmarks for M1 and the 5-cm method for DLPFC targeting. M1 (light blue; mean in dark blue), DLPFC without VR headset (light green; mean in dark green), and DLPFC with VR headset (light pink; mean in dark pink). **(D)** Coil center positions at the F3 location of the 10-10 EEG system. F3 without VR headset (green), and F3 with VR headset (light pink; mean in dark pink). Wearing a VR headset increased coil-to-scalp distance at DLPFC targets by 32.5 ± 4.4 mm (mean ± SD; coil-to-scalp distance displacement).

The M1 motor spot was first identified using MNI coordinates (Figure 1B), and the coil was then moved 5 cm anteriorly to target the DLPFC according to a commonly used clinical approach. Across all trials (5 per model, 25 total), the Meta Quest 2 (MQ2) headset prevented positioning at this target under standard coil-to-scalp distance constraints (Figure 1B). Comparable results were obtained when M1 was defined using anatomical landmarks before shifting 5 cm anteriorly (Figure 1C).

We next assessed direct prefrontal targeting using the F3 position of the 10-10 EEG system, approaching the target from above rather than anteriorly in the same coil orientation (Figure 1D). In this configuration, the VR headset did not prevent placement of the coil center over F3, but the headset positioned between the coil and the scalp markedly increased coil-to-scalp distance from the 4 mm reference position to 36.5 ± 4.4 mm (mean ± SD), which corresponded to an average increase of 32.5 ± 4.4 mm. Together, these results show that VR headsets constrain DLPFC targeting.

### Systematic assessment of neocortical targeting across coil orientations

To systematically assess neocortex accessibility, we used the 10-10 EEG system, which provides broad coverage with 30 positions per hemisphere and 9 along the midline. Midline and extreme lateral positions (F9, FT9, TP9, P9) were excluded, as they are rarely targeted in TMS applications. The remaining 26 positions were evaluated using eight coil orientations per site, each rotated in 45° increments. Targeting constraints were quantified as the coil-to-scalp distance displacement between coil center positions obtained with and without a VR headset. Across positions, measurements were first averaged across orientations and then across subjects, yielding a mean coil-to-scalp distance displacement per region. Values were sorted, normalized to the range [0, 1] using min-max normalization, and visualized as a normalized topographical distribution (color-coded by mean coil-to-scalp distance displacement).

The 10 positions with the largest displacement are shown in Figure 2B. AF7 exhibited the largest mean displacement (47.4 ± 2.83 mm), followed by other prefrontal sites including F3 (31.1 ± 7.39 mm). Based on these findings, subsequent analysis focused on prefrontal regions. An orientation-resolved analysis revealed marked differences in coil displacement across angles (Figure 3). At F3, the largest displacement was observed at 225°, whereas the smallest displacement occurred at 135°, corresponding to posteriorly directed coil orientation toward the midline.

**Figure 2:**
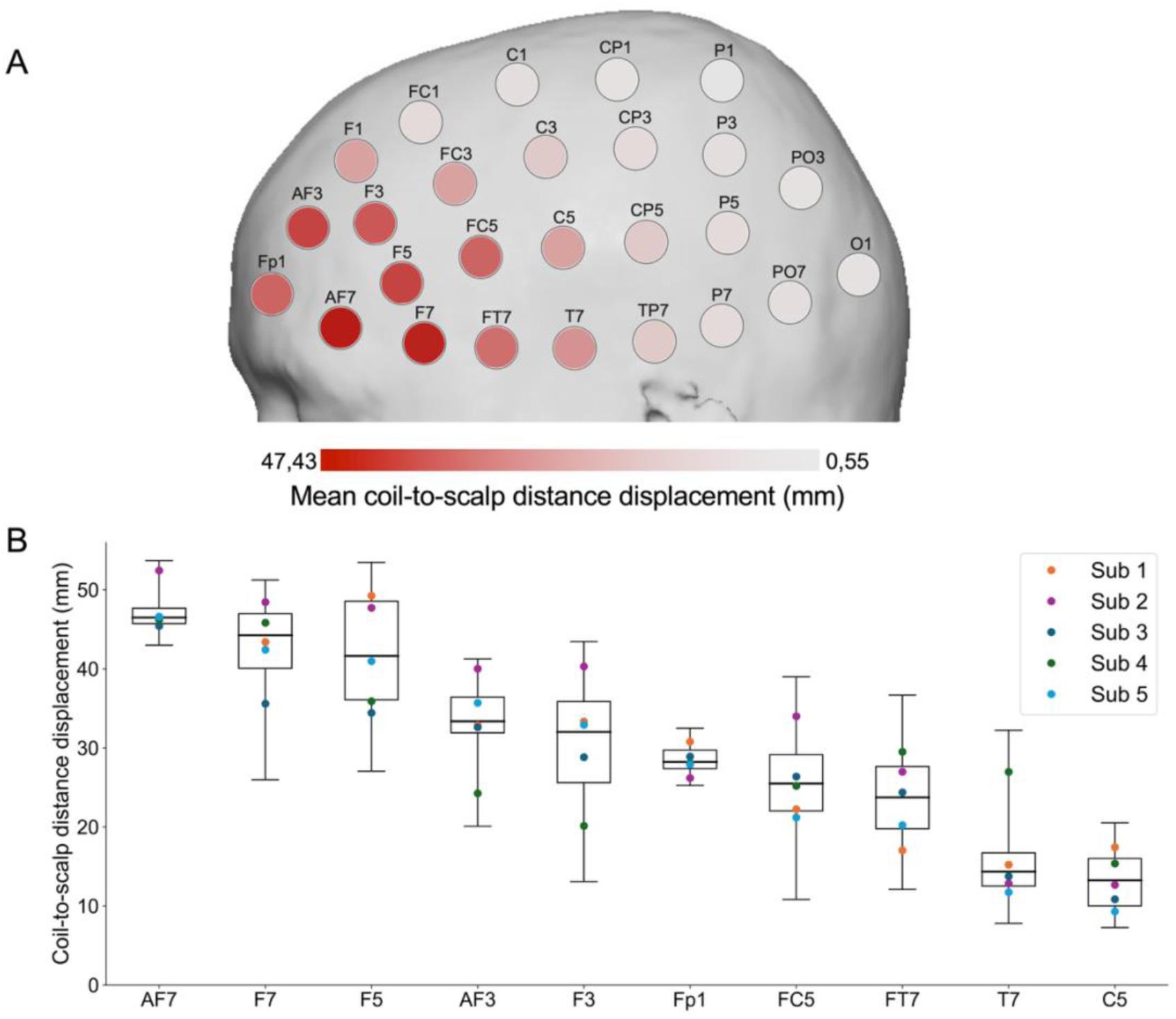
Neocortical targeting constraints across 10-10 EEG system positions. (**A)** Spatial distribution of mean coil-to-scalp distance displacement between intended and achievable coil positions across all subjects, min-max normalized and visualized as a color gradient (lighter shades indicate smaller displacement). (**B)** The 10 positions with the largest coil-to-scalp distance displacement. Box plots show distribution across positions and subjects (n=40; median, IQR, range); overlaid colored dots indicate mean subject-specific values.

**Figure 3:**
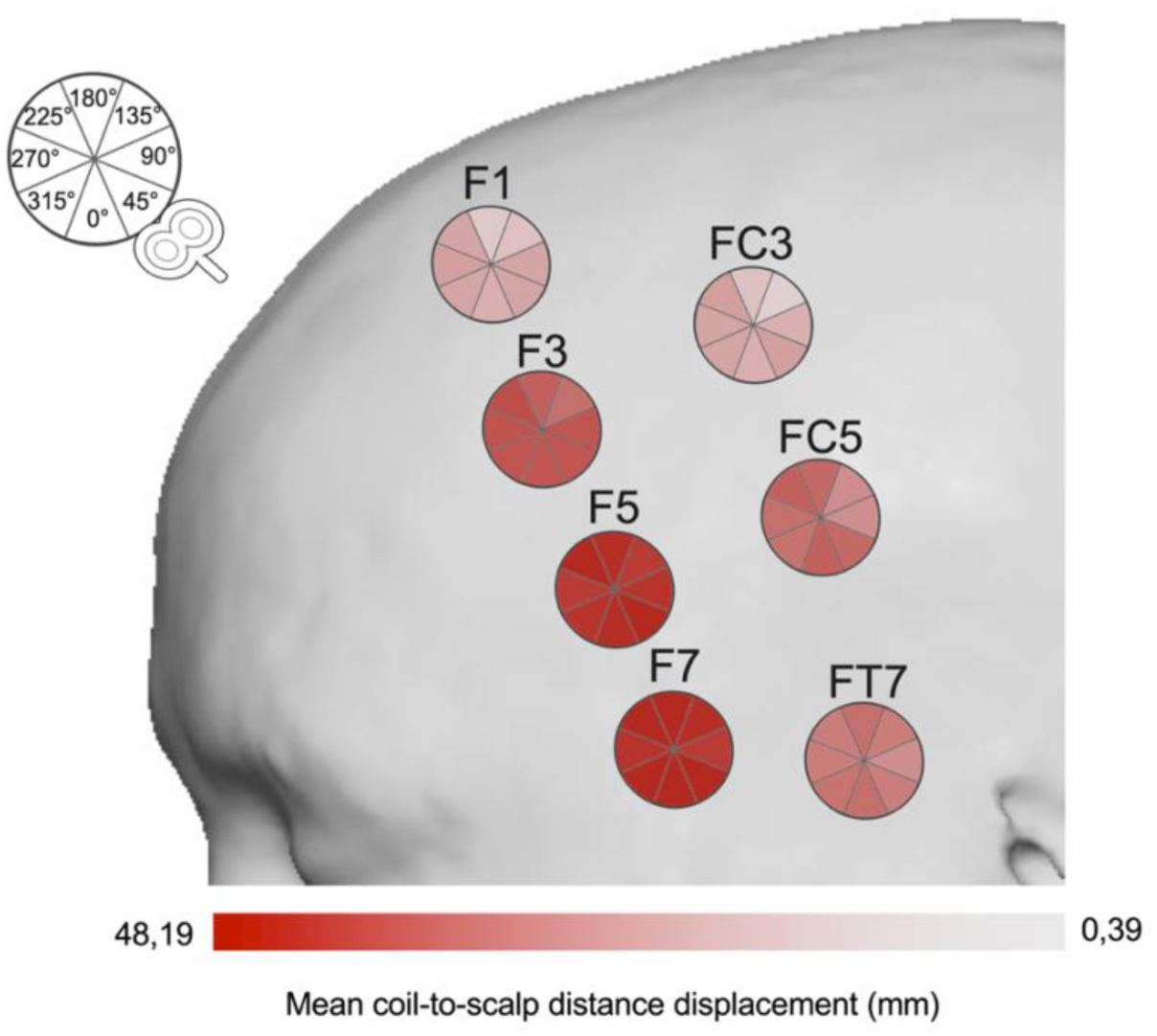
Orientation-dependent targeting constraints in prefrontal regions. Sector-based representation of orientations at selected prefrontal positions. Each sector corresponds to one of eight coil orientations (45° increments), with color indicating min-max normalized mean coil-to-scalp distance displacement across all subjects, lighter shades indicate smaller displacement. Min-max normalization was performed across the full dataset, including positions not shown.

### Comparison with a smaller VR headset

To assess whether headset geometry influences targeting constraints, measurements were repeated using a smaller headset. The Bigscreen Beyond (BB) VR headset (52.4 × 143.1 mm) is substantially more compact than the MQ2, with approximately half the width and one-quarter the height (Figure 4A; Table 1). Due to imperfect fit of the 3D-printed model in some head geometries, small gaps between the headset and the face were observed, which may have slightly increased coil-to-scalp distance in these cases. Despite this limitation, the smaller headset consistently reduced coil displacement over prefrontal regions. This effect was evident across orientations at F3, with significantly lower coil-to-scalp distances compared to the MQ2 (p ≤ 0.004 for all orientations) (Figure 4B, C).

**Figure 4:**
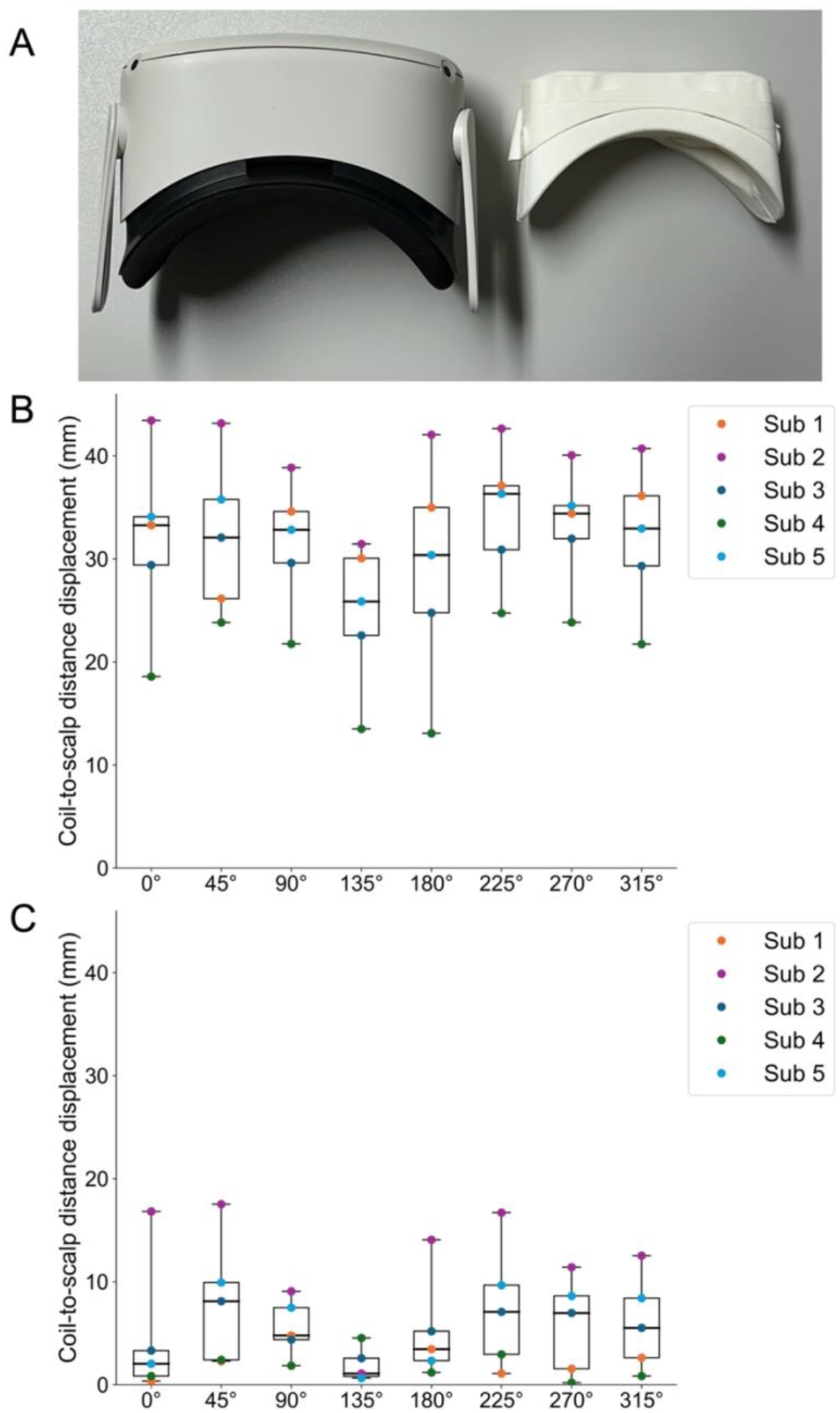
Comparison of prefrontal targeting constraints with different VR headsets. **(A)** Meta Quest 2 headset (left, without strap) and 3D-printed Bigscreen Beyond (right), shown in top view. **(B)** Coil-to-scalp distance displacement at F3 across coil orientations for the Meta Quest 2 headset. Box plots represent the distribution of values across subjects (n=5; median, IQR, range); overlaid colored dots indicate subject-specific values. **(C)** Coil-to-scalp distance displacement at F3 across coil orientations for the Bigscreen Beyond headset. Box plots represent the distribution of values across subjects (n=5; median, IQR, range); overlaid colored dots indicate subject-specific values.

### Adjusting stimulation intensity to achieve comparable E-fields in the target region

To systematically evaluate changes in E-field characteristics at the target site (F3), simulations were performed using the reference condition (1% MSO, 4 mm coil-to-scalp distance). Coil-to-scalp distance was then increased from 4 to 40 mm in 1 mm increments while keeping all other parameters constant. Both absolute E-field metrics and region-specific measures within the predefined region of interest were analyzed.

Across all subjects, increasing coil-to-scalp distance resulted in a progressive reduction in E-field strength and a concomitant increase in spatial spread. To maintain the reference E-field strength (P98, 98th percentile of the E-field magnitude), each additional millimeter of distance required an increase in stimulation intensity of approximately 4.5-4.7% MSO, consistent with values reported in individual-level E-field modeling studies (Caulfield et al., 2024). Changes in field spread were approximately linear, increasing ∼0.6% per millimeter, corresponding to +3.3% at 10 mm and +18.4% at 35 mm relative to the 4 mm baseline (Figure 5).

**Figure 5:**
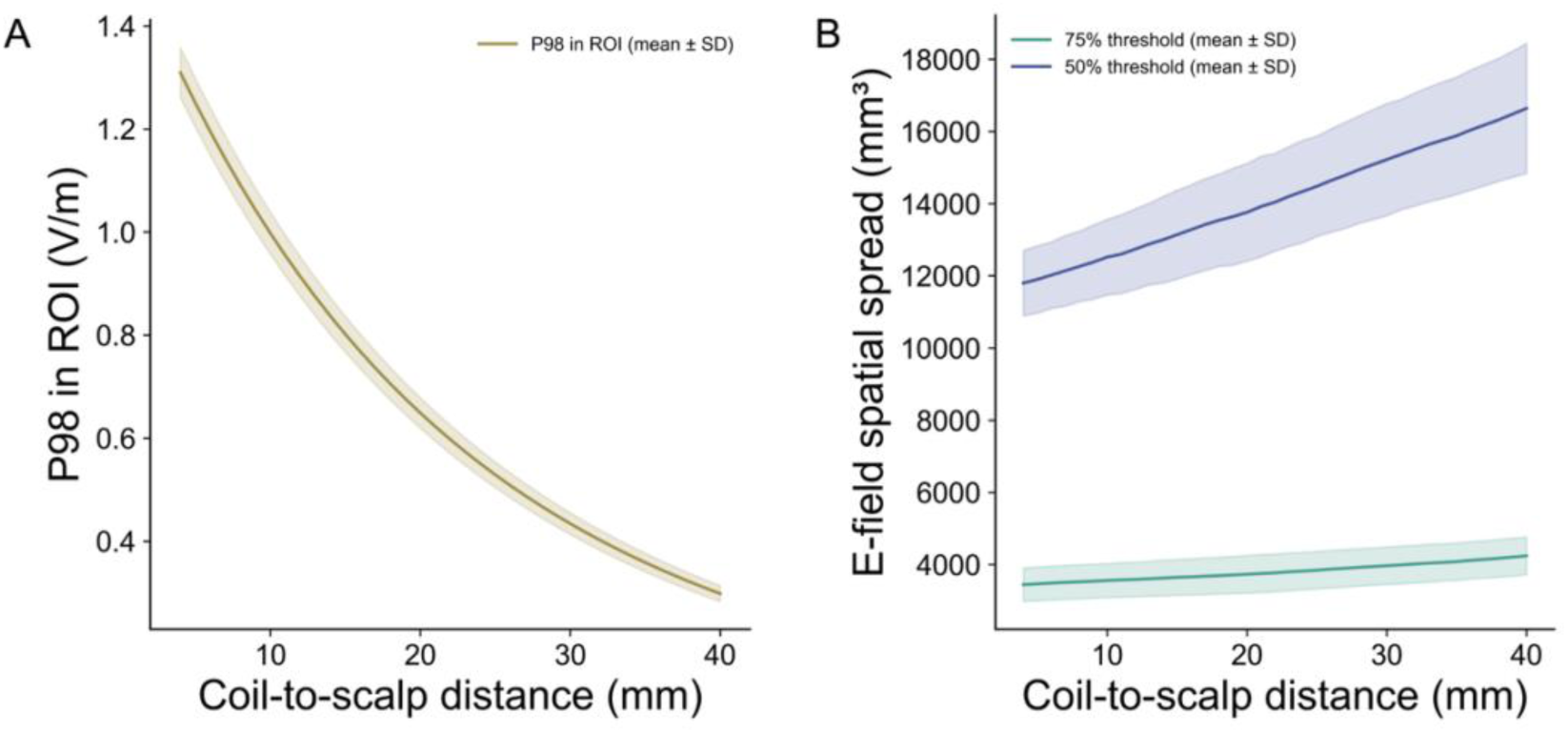
Effect of coil-to-scalp distance on E-field strength and spatial spread. **(A)** Change in E-field strength (98th percentile of the E-field magnitude) as a function of coil-to-scalp distance (mean ± SD). **(B)** Change in E-field spatial spread (defined as gray matter volume exceeding 75% and 50% of the peak E-field) as a function of coil-to-scalp distance (mean ± SD).

Of greater clinical relevance was whether the reduction in E-field strength could be compensated by increasing stimulator output. To address this, we calculated the intensity scaling factor required to restore reference field parameters (defined at 1% MSO, 4 mm coil-to-scalp distance, F3) across all simulated distances (Figure 6).

**Figure 6:**
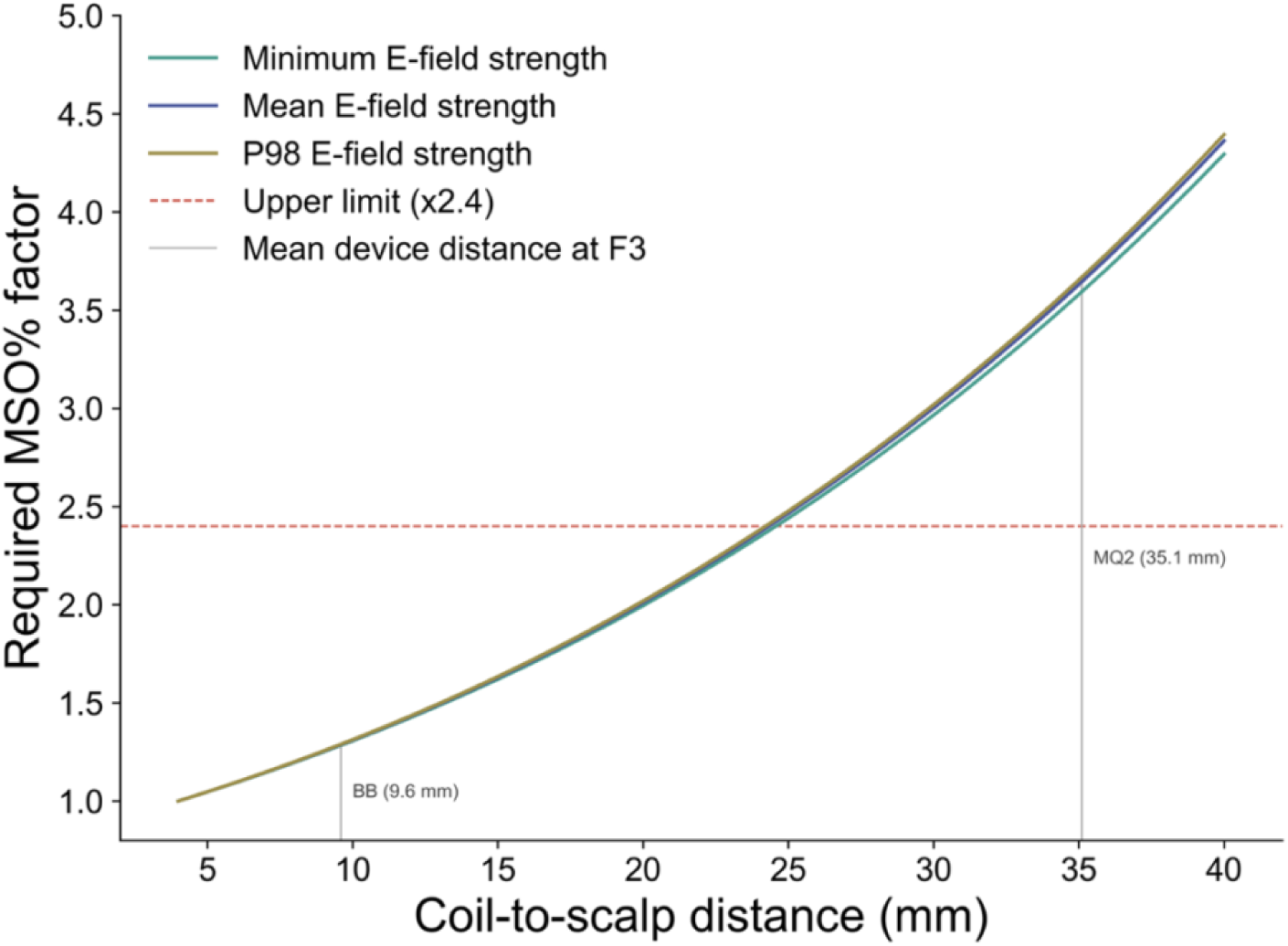
MSO scaling required to restore reference E-field. Required MSO scaling factor to match reference E-field parameters (minimum, mean, and P98) within F3 as a function of coil-to-scalp distance (mean ± SD), expressed relative to the reference condition (1% MSO, 4 mm). The horizontal dashed red line indicates the upper compensable range (scaling factor 2.4, corresponding to 100% MSO). Vertical lines denote mean coil-to-scalp distances (including the 4 mm reference offset) for the two VR headsets (Bigscreen Beyond: 9.6 mm; Meta Quest 2: 35.1 mm).

To define a clinically feasible compensation range, the experimental reference (1% MSO) was mapped to typical clinical stimulation intensities (42-62% MSO, depending on the device), yielding a maximum allowable scaling factor of 2.4 before reaching 100% MSO.

Restoration of target E-field parameters was feasible up to a coil-to-scalp distance of ∼23-25 mm (depending on the metric), corresponding to an additional displacement of ∼19-21 mm from the reference condition. This range encompasses the smaller VR headset (BB, mean displacement at F3: 5.6 mm; corresponding coil-to-scalp distance: 9.6 mm). In contrast, the larger headset (MQ2, mean displacement: 31.1 mm; corresponding coil-to-scalp distance: 35.1 mm) exceeded the compensable range across all metrics, indicating that field restoration is not achievable within maximum stimulator output limits (100%).

### Assessment of technical compatibility

To assess whether TMS interferes with VR headset functionality, we tested single-pulse stimulation (5-75% MSO in 10% increments and 100% MSO) and intermittent theta-burst stimulation (iTBS 600; 50% MSO). All experiments in this section were conducted using the MQ2 headset. During stimulation, a 4K video with dynamic audio was presented through the headset, and output was monitored via camera recording of the display (Figure 7).

**Figure 7:**
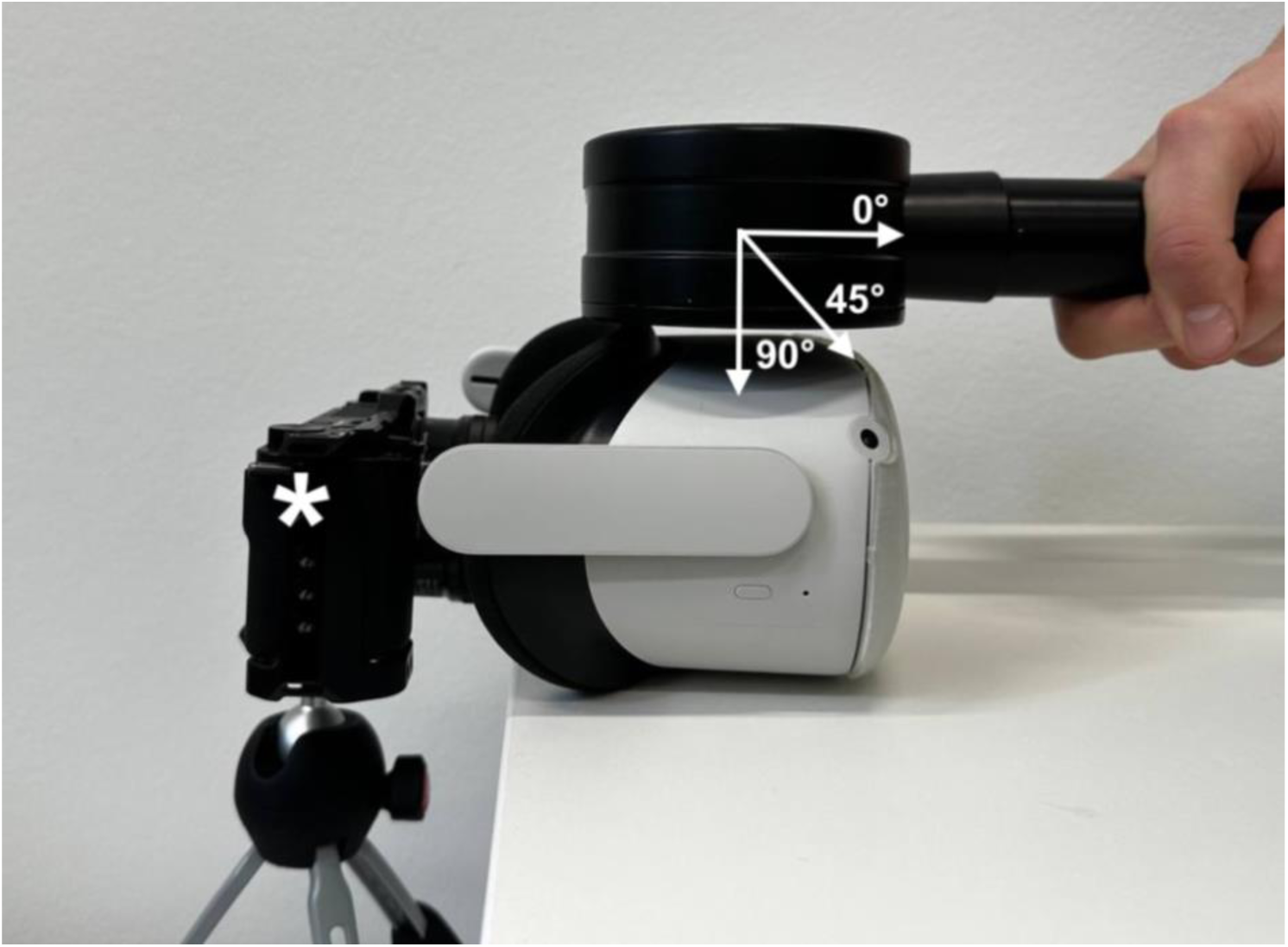
Assessment of technical compatibility during TMS. Photograph of the experimental setup. Single-pulse and iTBS protocols were applied to evaluate potential interference of TMS with VR headset functionality (Meta Quest 2). Visual and auditory output was monitored by recording the headset display using a camera (asterisk).

For single-pulse stimulation, the coil was positioned centrally over the headset at orientations of 0°, 45°, and 90°, as well as two lateral positions (2.5 cm medial to the left and right headset edges), approximating the positions of internal displays. In all conditions, the coil was placed in direct contact with the headset. No interference with visual or auditory output was observed. For iTBS 600, the coil was positioned centrally at 90° in direct contact with the headset. At 50% MSO, stimulation induced video interruption immediately after 600 pulses, accompanied by a transient overheating warning. Increasing the coil-to-headset distance to 35 mm delayed this effect, which then occurred after three consecutive iTBS 600 sessions (i.e., after 1800 pulses).

To assess interference under realistic targeting conditions, the coil was positioned over F3 tangentially to the scalp, with the handle oriented postero-laterally at 45°, while the headset was worn on the model. Three consecutive iTBS 600 sessions were applied, with 10-second intervals between sessions to allow manual monitoring for overheating warnings. Under these conditions, no overheating or functional interference was observed. Together, these findings indicate that while hardware interference is limited under realistic targeting conditions, headset geometry remains the primary constraint for effective TMS targeting.

## DISCUSSION

The integration of VR and TMS holds significant potential for advancing human neuroscience and clinical applications, particularly for studying and modulating neural processes in health and disease. Our findings demonstrate substantial constraints in targeting specific brain regions, such as the DLPFC, when using immersive VR headsets. These constraints can be addressed primarily through headset geometry adaptation, rather than stimulation parameter adjustment. Standard VR headsets, such as the Meta Quest 2 (Table 1), introduce technical constraints that substantially increase coil-to-scalp distance, with displacements at DLPFC sites reaching 31.1 mm at F3 and up to 47.4 mm at AF7, values that far exceed variations previously attributed to hairstyle or electrode caps (Caulfield et al., 2024), and that likely place effective prefrontal stimulation beyond the reach of standard clinical intensity scaling. These findings suggest that compact headset designs represent a critical consideration for future VR-TMS applications targeting the DLPFC.

The principal mechanism underlying these constraints is geometric: VR headsets occupy prefrontal space, and coil placement must accommodate their physical profile. Our systematic assessment using the 10-10 EEG system indicates that this constraint is not uniform across the scalp - displacement is concentrated over frontal and prefrontal regions, while parietal and central sites remain largely accessible. This topographic specificity has direct implications for target selection: VR-TMS applications targeting motor or parietal cortex appear feasible with current-generation headsets, whereas DLPFC stimulation, the dominant target in clinical rTMS for depression and cognitive modulation, is selectively compromised. Coil orientation further modulates these constraints: at F3, displacement was largest at 225° and smallest at 135°, indicating that orientation optimization may partially reduce, but not eliminate, headset-imposed displacement.

In exploring potential solutions, our simulation results show that adjusting stimulation intensity can partly compensate for increased coil-to-scalp distances, with defined limits. Restoring reference E-field parameters at F3 required approximately 4.5-4.7% additional MSO increase per millimeter of coil-to-scalp distance and was feasible up to approximately 23-25 mm displacement. For the Meta Quest 2, mean displacement of 31.1 mm exceeded the compensable range across all metrics, meaning restoration would require surpassing 100% MSO, which is not clinically feasible. Therefore, hardware geometry, rather than stimulation parameters, appears to be the primary constraint for standard-sized headsets.

The more compact Bigscreen Beyond headset demonstrates that these limitations are not intrinsic to VR as a technology, but are a function of headset form factor. Its substantially reduced frontal configuration brought mean F3 displacement to 5.6 mm, well within the compensable range, while control measurements with the physical device confirmed the geometric accuracy of the printed model. At this displacement, intensity compensation remained within clinical bounds with increases in spatial spread of approximately 3-4%, suggesting that compact headset designs can restore effective prefrontal stimulation with modest increases in spatial spread. This finding highlights the importance of prioritizing headset compactness as a primary engineering parameter in future VR-TMS development, rather than a secondary consideration. Modular or recessed frontal designs, already explored in the context of EEG-VR integration (Weber et al., 2021), offer a potential solution for resolving similar hardware conflicts without compromising VR immersion.

To assess whether TMS interferes with VR headset functionality, we performed an exploratory evaluation of technical compatibility under a range of stimulation conditions. Single-pulse stimulation produced no visual or auditory interference across intensities and orientations and iTBS applied with the coil positioned tangentially over F3 caused no overheating or functional disruption across three consecutive sessions. Interference was only observed when the coil was placed in direct contact with the headset surface during high-frequency protocols. While these findings are encouraging, they represent an initial assessment rather than a comprehensive characterization of electromagnetic compatibility and further systematic investigation under a broader range of conditions would be required before drawing conclusions.

Several limitations need consideration. This pilot study examined five individual head models and while intersubject variability in displacement was systematically characterized, generalization to broader anatomical variation, such as different age groups or pathological populations, requires further investigation. The Bigscreen Beyond headset was evaluated using a 3D-printed model and although control measurements showed no substantial deviations, minor geometric discrepancies may have slightly increased coil-to-scalp distance estimates in some subjects. Additionally, simulations assumed a linear MSO-to-E-field relationship and a fixed coil-current rate, consistent with standard modeling practice but abstracted from the full complexity of individual neuroanatomy and stimulator variability.

An important open question concerns the potential influence of the VR environment itself on cortical excitability. Immersive VR may modulate baseline brain states in ways that affect resting motor threshold and the stimulation intensities required to achieve target E-field effects - factors that were not addressed in the present study and that future work should examine.

In summary, while combining VR and TMS holds considerable potential, particularly for neurocognitive and psychiatric interventions, technical challenges currently limit the accessibility of critical brain regions with standard hardware. By establishing quantitative boundaries for compensable and non-compensable coil-to-scalp displacement, these findings provide a practical framework for researchers and clinicians designing concurrent VR-TMS protocols and highlight the hardware design requirements that future integrated systems will have to meet.

## AUTHOR CONTRIBUTIONS

FA: Conceptualization, methodology, software, formal analysis, investigation, data curation, visualization, writing - original draft, writing - review and editing; PH: conceptualization, investigation, writing - review and editing; ZT: methodology, validation, software, writing - review and editing; AV: conceptualization, methodology, resources, writing - original draft, writing - review and editing, supervision, funding acquisition.

## DATA AND CODE AVAILABILITY STATEMENT

The data and analysis scripts that support the findings of this study are available from the corresponding author upon reasonable request.

## Acknowledgements

This work was supported by the Federal Ministry of Education and Research, Germany (BMBF, 01GQ2205A to AV).

